# Metagenomic peek into a corn mummy

**DOI:** 10.1101/2024.07.02.601727

**Authors:** Norbert Solymosi, Bernadett Pap, Sára Ágnes Nagy, Adrienn Gréta Tóth, Flóra Judit Kevély, Gergely Maróti, István Csabai, Katalin Kóthay, Donát Magyar

## Abstract

Numerous studies have shown that metagenomics has opened a dimension in reading the contents of archaeological remains as time capsules. Corn mummies are ritual objects from ancient Egypt, created by forming human-shaped figures from cereal grains grown in a mixture of water and earth. The aim of our study was to determine whether ancient DNA could be preserved in the mummy, and if so, which organisms it might have originated from. To find answers, we performed metagenomic analyses on samples taken from a corn mummy dating to the second half of the third century BC. Alongside a number of clearly modern contaminants, we identified organisms that cannot be excluded as being of historical origin. Besides considerable amounts of bacterial sequences belonging to the genus *Bacillus, Mesobacillus, Metabacillus, Neobacillus, Niallia, Peribacillus* and *Paenibacillus*, we also found traces of plants, animals, and humans. Sequences assigned to the genus *Triticum* showed the highest similarity to ancient *T. turgidum* ssp. *dicoccum* specimens from Egypt and the southern Levant. The fragments identified as of Lepidopteran origin showed the greatest similarity to Sphingidae genomes. Analysis of the human-derived sequences revealed L3 (mtDNA), E, and J (Y chromosome) haplotypes, which are common lineages in Africa today.

## Introduction

Metagenomics, based on next-generation sequencing, yields further insights into complex communities and ecosystems. In addition to recent, contemporary samples, this approach also allows the detection of traces of organisms that once lived and were trapped in a sample as a time capsule based on the DNA content of archaeological remains.^1–4^ Metagenomic tools have been used to identify the genome of the oldest known *Mycobacterium leprae* from the bones of an Egyptian human mummy.^5^ Khairat et al.^6^ also detected microbial and plant genome fragments in samples from human mummies. A special type of mummy is the corn mummy, which was made in Egypt between the 3rd century BC and the 1st century AD as part of the religious rites of the Khoiak festival. In the ceremony, the grain was grown in a mixture of 1 hin (ancient unit of volume, ∼0.47 L^7^) of barley and/or wheat,^8,9^ 1 hin of water, and 4 hin of sand for 9 days (between 12 and 21 of the month of Khoiak, around 22-31 December), and it was watered by river water.^10^ Then, the formed mummy was placed on dried myrrh, embalmed, painted, wrapped in papyrus and clothed with textiles, and finally placed in a sycamore coffin.^11^ The results of metagenomic research on ancient Mesopotamian clay bricks by Arbøll et al.^12^ inspired us to conduct a similar study on a corn mummy. Our study aimed to determine whether ancient DNA could be preserved in the mummy and, if so, which organisms it might have come from. To this end, metagenomic analysis was performed on samples from one of the two corn mummies in the Egyptian Collection of the Museum of Fine Arts, Budapest, Hungary.

## Material and methods

### Sampling

The Egyptian Collection of the Budapest Museum of Fine Arts holds two corn mummies. One of them (inv. no. 60.22-E) is completely intact and has a sarcophagus, while the other (inv. no. 68.2-E) is without a sarcophagus and is broken crosswise (Figure 1). Object no. 68.2-E was acquired by the Museum in 1968. Experts estimate it is from Central Egypt and date it to the second half of the 3rd century BC on stylistic grounds.^11^ The object from which the sample was taken is not on exhibition; it is stored under special conditions in a closed museum storage facility, where the sampling also took place. Sampling permit was granted by the Egyptian Collection of the Museum of Fine Arts.

**Figure 1.**
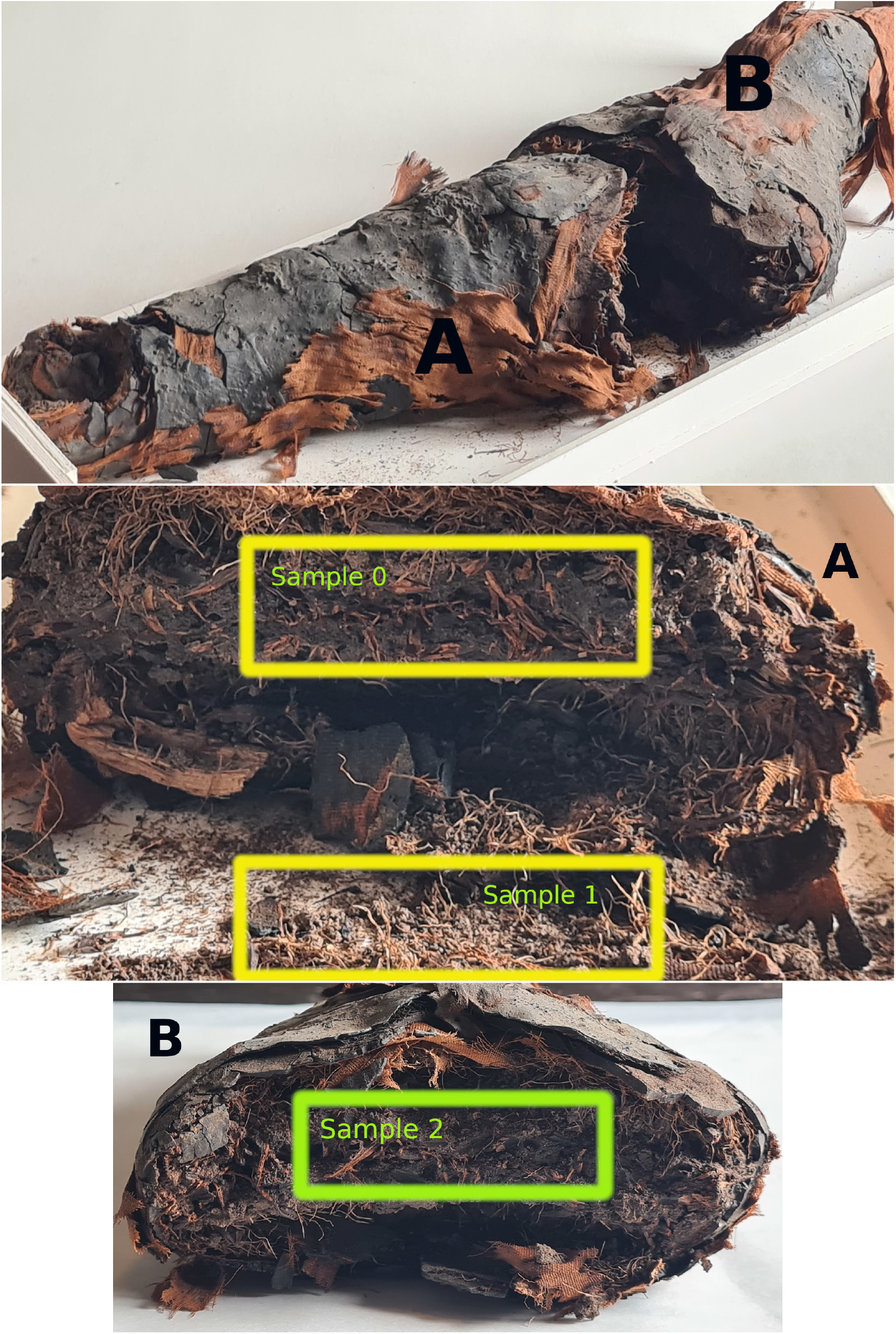
Corn mummy 68.2-E sampled. The top image shows the whole mummy, the middle image shows part A with sampling areas 0 and 1, and the bottom image shows part B with sampling area 2.

Due to the high value and rarity of the specimen, we took samples from the fractured mummy (68.2-E) with great care, disturbing only the fracture surface. On the very first occasion (Sample 0), specimens were taken from the surface shown in

Figure 1 by “thin needle biopsy” (Table 1). This was done by inserting a sterile, long blood collection needle deep (1-2 cm) into part A of the mummy and withdrawing the needle with a syringe to create a vacuum. The remains in the needle lumen were then collected into a sterile tube. For Sample 1, we used the fragments that had fallen off the mummy (Figure 1). For Sample 2, using a sterile syringe cone, we scraped 1-2 mm of the fracture surface from part B of the split mummy. Continuing this procedure, a sample was taken from the material deeper than the surface using a sampling tube. The collected samples were transported to the sequencing lab in a sterile, closed tube.

**Table 1.** Summary of sampling and sequencing details. For library preparation the NEBNext® Ultra™ II DNA Library Prep Kit for Illumina (Cat.Num.: E7645L) was used. Sequencing was performed at Seqomics Ltd. by NextSeq sequencer using TG NextSeq® 500/550 High Output Kit v2 (300 cycles). The sampling positions of the corn mummy are shown in Figure 1.

| ID | Sampling | Depth | Extraction protocol |
| --- | --- | --- | --- |
| Sample 0 | thin needle biopsy | 1-2 cm | Kistler L.<sup>13</sup> |
| Sample 1 | fallen fragments | surface | in-house developed special method |
| Sample 2 | syringe cone | 1-2 mm | Latorre et al.<sup>14</sup> |

### DNA extraction, library preparation and sequencing

Samples from the corn mummies were delivered to HUN-REN Biological Research Centre (BRC) for DNA extraction. A separated laboratory equipped with HEPA filters and built-in UVC lamps (3x 36 W, radiating at 254 nm wavelength, automatically operating every day between 5 and 6 am) was used for all aDNA extractions. Dedicated tools (pipettes, centrifuges, incubators, glassware, and plasticware) and chemicals were used for aDNA purification to prevent contamination. Extraction blank samples to control possible contaminations were included in all cases. An Agilent TapeStation 4150 system was used for the QC of the extracted DNA from all samples.

Kistler’s CTAB Protocol^13^ was used to extract aDNA from 200 mg material of Sample 0. For Sample 1, cell pellets were resuspended in 567 *µ*L of 1xTE (10 mM Tris, 1 mM EDTA pH 8.0) buffer (pH 8.0) containing 2 mg/mL of lysozyme and incubated for 30 minutes at 37°C. After adding 30 *µ*L 10% sodium dodecyl sulfate (SDS) and 3 *µ*L 20 mg/mL Proteinase K incubated for 1 hour at 37°C. Samples were then incubated for 20 minutes at 65°C with 150 *µ*L of 5 M NaCl and 80 *µ*L of CTAB/NaCl solution (700 mM NaCl/10% CTAB). Following incubation, extracts were purified using phenol-chloroform extraction, and aDNA was recovered by isopropanol precipitation. Pelleted aDNA was washed with 70% ethanol, allowed air dry, and resuspended in 32 *µ*L of 1xTE buffer. In the case of Sample 2, following the Basic Protocol 1 published by Latorre et al.,^14^ 20 and 200 mg samples were used to isolate aDNA.

Paired-end libraries were prepared at HUN-REN BRC using the NEBNext® Ultra™ II DNA Library Prep Kit for Illumina (Cat.Num.: E7645L). Sequencing was performed at Seqomics Ltd. genomic center (Mórahalom, Hungary) in a quality-controlled NGS platform, special care was taken with the libraries prepared from aDNA (libraries were not pooled for sequencing with libraries from other sources). Paired-end fragment reads were generated for the aDNA libraries on an Illumina NextSeq sequencer using TG NextSeq® 500/550 High Output Kit v2 (300 cycles). Primary data analysis (base-calling) was performed using the Bbcl2fastq software (v2.17.1.14, Illumina).

### Bioinformatics

The raw paired-end FASTQ files were merged, trimmed and filtered by AdapterRemoval (v3.0.0-alpha2, `--trim-ns--trim-qualities --trim-min-quality 20 --min-length 30 --min-complexity 0.3`).^15^ The remaining reads were taxonomically classified by Kraken2 (v2.1.3, `--confidence 0.2`)^16^ on the NCBI Core NT^17^ database (built: 28/12/2024, available at https://genome-idx.s3.amazonaws.com/kraken/k2_core_nt_20241228.tar.gz). The taxon abundance results were adjusted by Bracken (v3.1).^18^ The de novo assembly was performed by MEGAHIT (v1.2.9)^19^ with default settings. The assembled contigs to check the most probable origin were aligned by local BLAST^20^ on NCBI Core NT database (updated: 11/2/2025). For all contig and reference genome based aligment BWA (v0.7.17-r1188)^21^ was used, in the case of ancient samples with the settings: `aln -n 0.01 -l 16500 -o 2`^22^. The BAM files used for variant and consensus sequence calling, DNA damage analysis, contaminant analysis, and karyotype analysis were duplicated by Picard (v2.26.6, https://broadinstitute.github.io/picard/).

Post-mortem DNA damage was analyzed by estimating the frequency of C>T and G>A misincorporation using MapDamage (v2.3.0a0).^23^ The alignments used for this purpose were produced on the de novo assembled contigs and reference genomes of the most abundant species of genera with at least 1% relative abundance.

The phylogenetic analysis of present and ancient samples of the genus *Triticum* and *Hordeum* was based on the longest contig of the genus *Triticum*, which we generated by de novo assembly. For this purpose, we have downloaded paired-end Illumina-sequenced datasets of ancient and present *Triticum* and *Hordeum* samples (Table 2) from the NCBI Sequence Read Archive (SRA). The raw short reads from these external samples were preprocessed as described above. The ancient origin samples’ reads were mapped with the above-detailed BWA aln settings. Following the approach of Lev-Mirom et al.^24^, the reads from the downloaded present samples were mapped to our longest assembled contig using BWA-MEM. From the generated BAM files, the variant call and the consensus sequence call were performed using BCFtools (v1.14). For the phylogenetic analysis, we used consensus sequences with an N ratio of less than 5The gene-tree was constructed^25^ based on multiple sequence alignment by MAFFT (v7.490, `--auto`).^26^ The best substitution model was selected by functions of phangorn (v2.12.1) package^27^ based on the Bayesian information criterion (BIC). The generated neighbor-joining tree was optimized using the maximum-likelihood method. Bootstrap values were produced by 100 iterations. All data processing and plotting were done in the R environment (v4.6.0).^28^

**Table 2.** The *Triticum* and *Hordeum* external samples were used in phylogenetic analysis by the longest wheat origin contig. The short read data sets of the samples were downloaded from NCBI SRA, and aligned to the PV405787 sequence. The contigs generated from alignments with less than 5% missing sites were used for multiple sequence alignment and phylogenetic analysis (Figure 6). The ancient sample Tt_UC10164 is dated to 1130–1000 BC,^29^ while southern Levant samples obtained from the BioProject PRJEB64853 are dated to 4500-2400 BC.^24^

| ID | Biosample | Origin | Species | Alignment statistics |  |  |
| --- | --- | --- | --- | --- | --- | --- |
|  |  |  |  | Coverage % | Mean depth | Variable site count |
| *Ancient* |  |  |  |  |  |  |
| Tt_JK2283 | SAMEA114240246 | southern Levant | T. turgidum ssp. dicoccum | 100.0 | 72.0 | 57 |
| Tt_JK3015 | SAMEA114240247 | southern Levant | T. turgidum ssp. dicoccum | 100.0 | 122.5 | 56 |
| Tt_TU1126 | SAMEA114240248 | southern Levant | T. turgidum ssp. dicoccum | 99.3 | 52.9 | 73 |
| Tt_TU1128 | SAMEA114240249 | southern Levant | T. turgidum ssp. dicoccum | 93.3 | 25.4 |  |
| Tt_TU1129 | SAMEA114240250 | southern Levant | T. turgidum ssp. dicoccum | 100.0 | 112.2 | 49 |
| Tt_TU1130 | SAMEA114240251 | southern Levant | T. turgidum ssp. dicoccum | 100.0 | 89.8 | 60 |
| Tt_TU1131 | SAMEA114240252 | southern Levant | T. turgidum ssp. dicoccum | 99.4 | 37.4 | 76 |
| Tt_TU1145 | SAMEA114240253 | southern Levant | T. turgidum ssp. dicoccum | 100.0 | 75.7 | 76 |
| Tt_TU1146 | SAMEA114240254 | southern Levant | T. turgidum ssp. dicoccum | 100.0 | 68.2 | 66 |
| Tt_TU1147 | SAMEA114240255 | southern Levant | T. turgidum ssp. dicoccum | 100.0 | 64.4 | 71 |
| Tt_TU1148 | SAMEA114240256 | southern Levant | T. turgidum ssp. dicoccum | 98.4 | 36.6 | 71 |
| Tt_TU1149 | SAMEA114240257 | southern Levant | T. turgidum ssp. dicoccum | 100.0 | 56.4 | 78 |
| Tt_TU1150 | SAMEA114240258 | southern Levant | T. turgidum ssp. dicoccum | 100.0 | 110.3 | 56 |
| Tt_TU686 | SAMEA114240259 | southern Levant | T. turgidum ssp. dicoccum | 92.3 | 25.5 |  |
| Tt_TU692 | SAMEA114240260 | southern Levant | T. turgidum ssp. dicoccum | 45.3 | 1.8 |  |
| Tt_UC10164 | SAMEA5328956 | Egypt | T. turgidum ssp. dicoccum | 100.0 | 189.3 | 40 |
| *Present* |  |  |  |  |  |  |
| H_vulgare | SAMN40712268 | Sweden | H. vulgare | 76.9 | 5203.7 |  |
| Ta_23896 | SAMN13640692 | Turkey | T. aestivum | 100.0 | 34461.0 | 65 |
| Ta_23909 | SAMN13640693 | Morocco | T. aestivum | 100.0 | 37011.5 | 64 |
| Ta_23923 | SAMN13640694 | Ethiopia | T. aestivum | 100.0 | 40149.5 | 59 |
| Ta_24056 | SAMN13640700 | Turkey | T. aestivum | 100.0 | 35433.2 | 68 |
| Ta_24184 | SAMN13640701 | Israel | T. aestivum | 100.0 | 38297.6 | 66 |
| Ta_AZTEC | SAMN13640786 | France | T. aestivum | 100.0 | 36584.8 | 65 |
| Ta_Altgold | SAMN13640588 | Germany | T. aestivum | 100.0 | 41040.8 | 67 |
| Ta_Apache | SAMN13640593 | France | T. aestivum | 100.0 | 39808.7 | 65 |
| Ta_Hubel | SAMN13640590 | Germany | T. aestivum | 100.0 | 39057.1 | 64 |
| Ta_IG21134349 | SAMN13640756 | Turkey | T. aestivum | 100.0 | 38416.3 | 64 |
| Ta_IG21134447 | SAMN13640757 | Jordan | T. aestivum | 100.0 | 38705.0 | 64 |
| Ta_IG21134470 | SAMN13640758 | Egypt | T. aestivum | 100.0 | 37703.9 | 59 |
| Ta_PI345242 | SAMN20819193 | Macedonia | T. monococcum | 89.3 | 1068.0 |  |
| Tm_PI427645 | SAMN16131686 | ? | T. monococcum | 100.0 | 5953.6 | 133 |
| Tm_TA759 | SAMEA13856358 | Saudi Arabia | T. monococcum | 90.8 | 17150.9 |  |
| Tt_DEW_PI_196100 | SAMEA112910807 | ? | T. turgidum ssp. dicoccum | 100.0 | 8953.8 | 82 |

To estimate the contamination of human genome originated fragments, reads were aligned to mitochondria, and chromosome X sequences. The reads aligned to NC_012920.1 sequence (Revised Cambridge Reference Sequence, rCRS) were analysed by Haplocheck (v1.3.3)^30^ and ContamMix (v1.0-11).^31^ By ContamMix 311 worldwide mitogenomes^32^ were compared to our mitochondrial consensus sequence using multiple sequence alignment with MAFFT. The schmutzi (v1.5.7)^33^ applied contamination analysis was performed on the Reconstructed Sapiens Reference Sequence (RSRS) alignment. For the X chromosome contamination analysis ANGSD (v0.940, `X:5000000-154900000`, `-doCounts 1 -iCounts 1 -minMapQ 30 -minQ 20`)^34^ package, based on NC_000023.10 (hg19) alignment was used. Karyotype assignment was done by alignment on X and Y chromosomes (hs37d5) with a minimum mapping quality of 30.^35^ For the mtDNA haplogroup classification using the RSRS-based alignment, the mitochondrial consensus sequence is called by schmutzi, and assigned by Haplogrep3 (v3.2.2, metric: Kulczyński, PhyloTree Build: 17).^**?**^ The Y chromosome-based haplogroup prediction was performed by pathPhynder^36^ and yHaplo (v2.1.12.dev9+geb252e7, `--anc_stop_thresh 10`)^37^, using the hs37d5 genomes. The VCF input of yHaplo was called by BCFtools (v1.14, `--min-MQ 30 --min-BQ 30 --ploidy 1`).^38^

## Results

In the case of Sample 0, it was not possible to isolate enough ancient DNA for sequencing. The number of cleaned, merged reads was 2,933,736 in the Sample 1 sample and 3,506,737 in the Sample 2 sample. The length of the contigs generated by de novo assembly was longer in Sample 1 (n: 12,479, median: 482, min: 201, max: 540,009, IQR: 453) than in Sample 2 (n: 12,851, median: 384, min: 257, max: 8,314, IQR: 147). The extraction blank samples resulted in no detectable DNA. The length distributions of the reads realigned on these contigs, and the misincorporation pattern are shown in Figure 2. While the read length distribution of Sample 2 has a peak at 84 bp, that of Sample 1 at 170. The contig and read lengths distribution with the DNA damage pattern shown in Figure 2 suggests that Sample 1 is rather a modern contamination than historical DNA. The length distribution and misincorporation pattern of Sample 2 suggest its ancient-DNA origin. Considering this, only the analysis results of Sample 2 are presented below.

**Figure 2.**
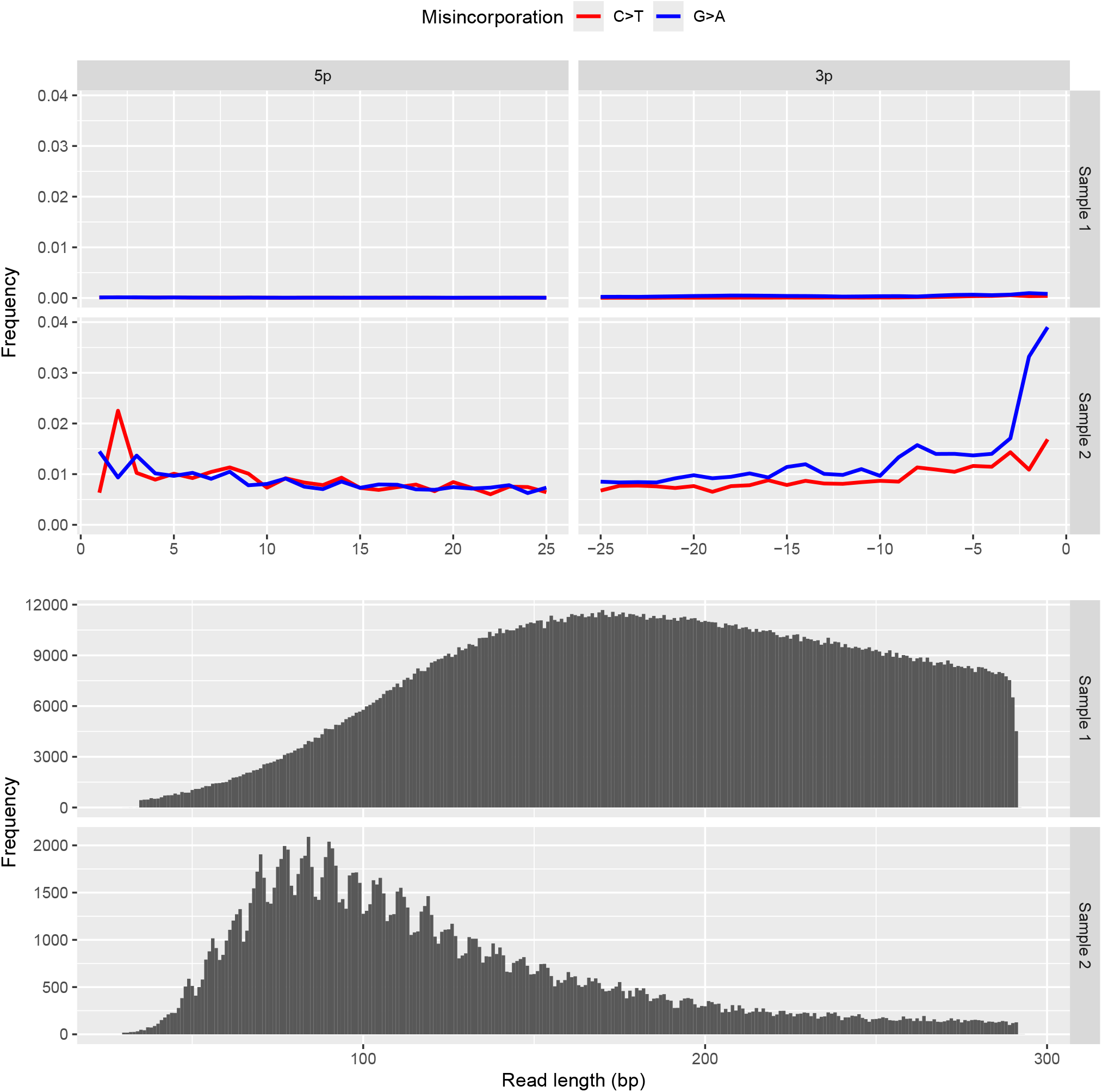
Misincorporation and read length plots. The number of aligned reads on the de novo assembled contigs was 2,010,050 and 170,184 in Samples 1 and 2, respectively.

Taxon classification of Sample 2 with 0.2 confidence resulted in 57.9% of hits being bacteria, 41.6% eukaryote, and the remaining viruses and archaea below 1%. Within the Bacteria, Bacilliota accounted for 83.2%, Actinomycetota for 7.7%, Pseudomonadota for 6.5%, and Bacteroidota for 2.3%. Within the Eukaryota, 88.2% are Chordata, 7.8% Streptophyta, and 2.7% Arthropoda. Within Chordata, 99.2% of the reads were classified to the *Homo* genus. In the phylum of Streptophyta, the genera above 1% with relative abundance were the *Triticum* (80.7%), *Populus* (7.8%), and *Aegilops* (5.1%). The genera with at least 1% relative abundance in the Arthropoda phylum were the *Bombyx* (91.3%), *Apis* (1.9%), *Ixodes* (1.5%), *Medioppia* (1.3%), *Drosophila* (1.1%), and *Oppiella* (1.1%). The genera with a minimal 1% relative abundance of classified reads were *Homo* (36.6%), *Niallia* (19.3%), *Bacillus* (8.1%), *Metabacillus* (8%), *Triticum* (2.6%), *Peribacillus* (2.3%), *Neobacillus* (2.2%), *Paenibacillus* (1.8%), *Cutibacterium* (1.7%), *Mesobacillus* (1.4%), *Bombyx* (1%). Within these genera, the most abundant taxa identified at the species level were *Homo sapiens, Niallia circulans, Bacillus paralicheniformis, Metabacillus* sp. KUDC1714, *Triticum aestivum, Cutibacterium acnes, Peribacillus frigoritolerans, Mesobacillus maritimus, Bombyx mori, Neobacillus* sp. DY30, *Paenibacillus* sp. FSL R5-0766. The DNA damage and length distribution patterns of reads aligned on the reference genomes of these species are shown in Figure 3-5.

**Figure 3.**
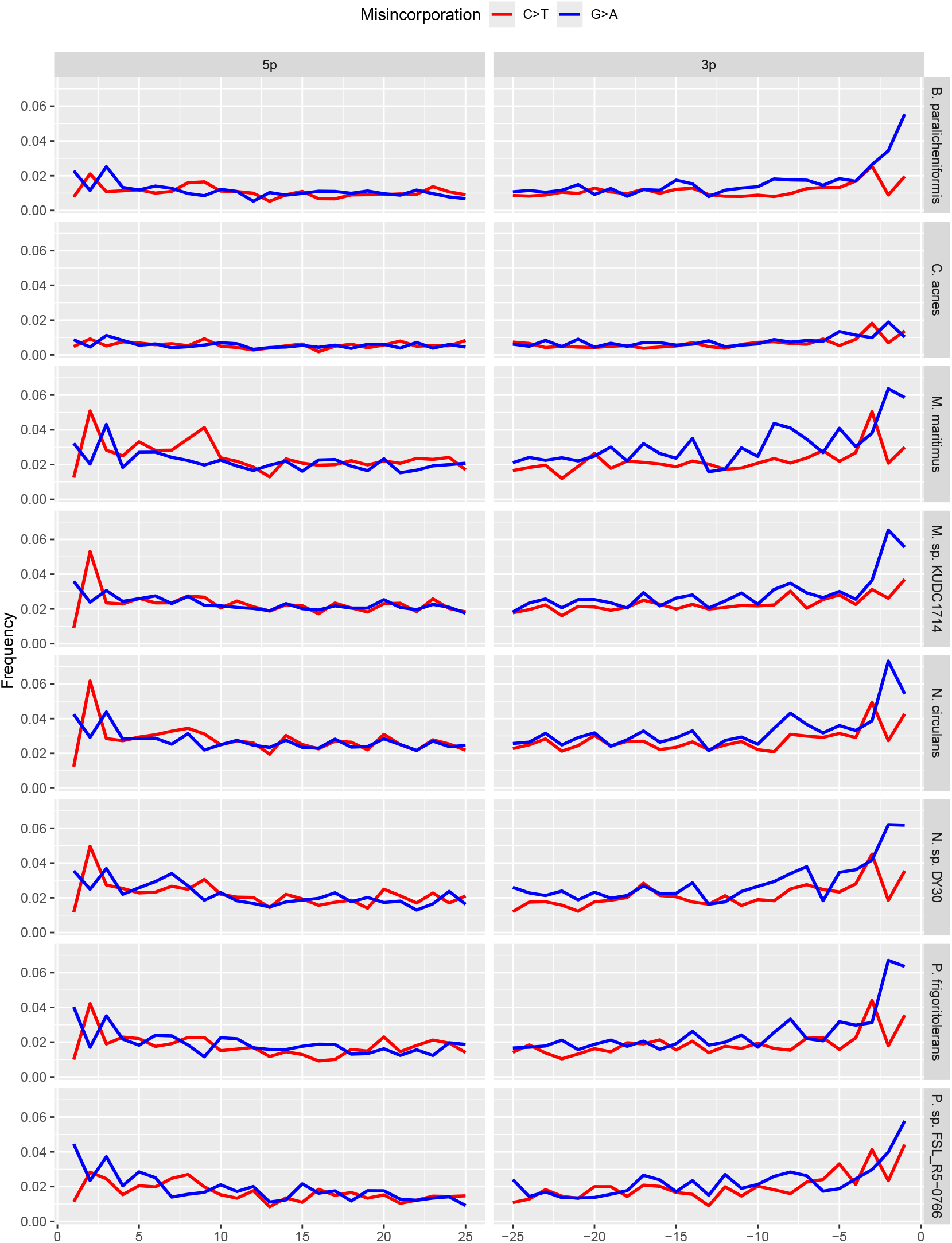
Damage plots of representative bacteria of the genera with at least 1% relative abundance. The genomes were used *Bacillus paralicheniformis* (GCF_002993925.1), *Cutibacterium acnes* (GCF_006739385.1), *Mesobacillus maritimus* (GCF_044803185.1), *Metabacillus* sp. KUDC1714 (GCF_014217835.1), *Neobacillus* sp. DY30 (GCF_030123065.1), *Niallia circulans* (GCF_013267435.1), *Peribacillus frigoritolerans* (GCF_022394675.1), *Paenibacillus* sp. FSL_R5-0766 (GCF_037971845.1).

**Figure 4.**
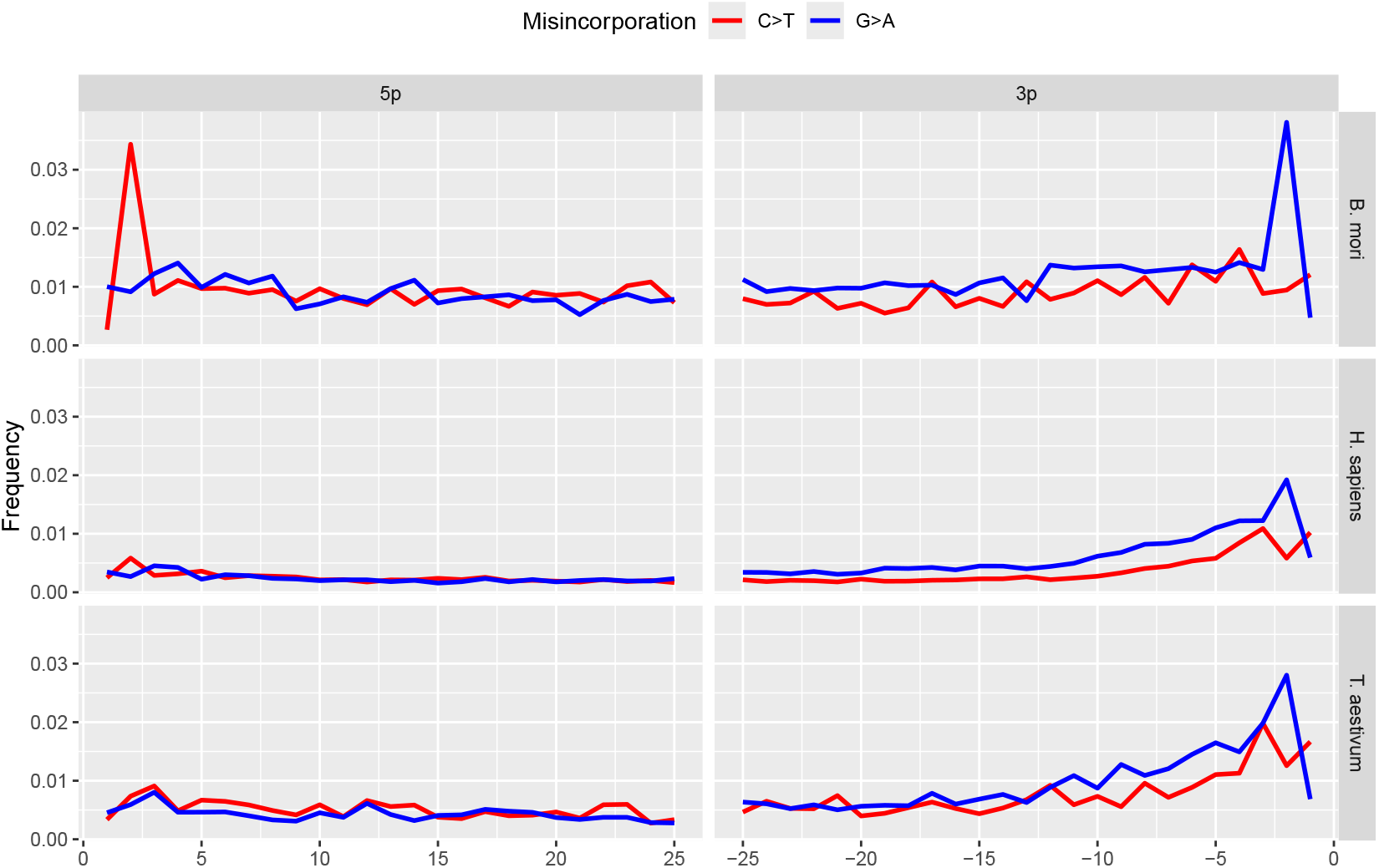
Damage plots of representative eukaryote species of the genera with at least 1% relative abundance. The genomes were used *Bombyx mori* (GCF_030269925.1), *Homo sapiens* (hs37d5), and *Triticum aestivum* (GCF_018294505.1).

**Figure 5.**
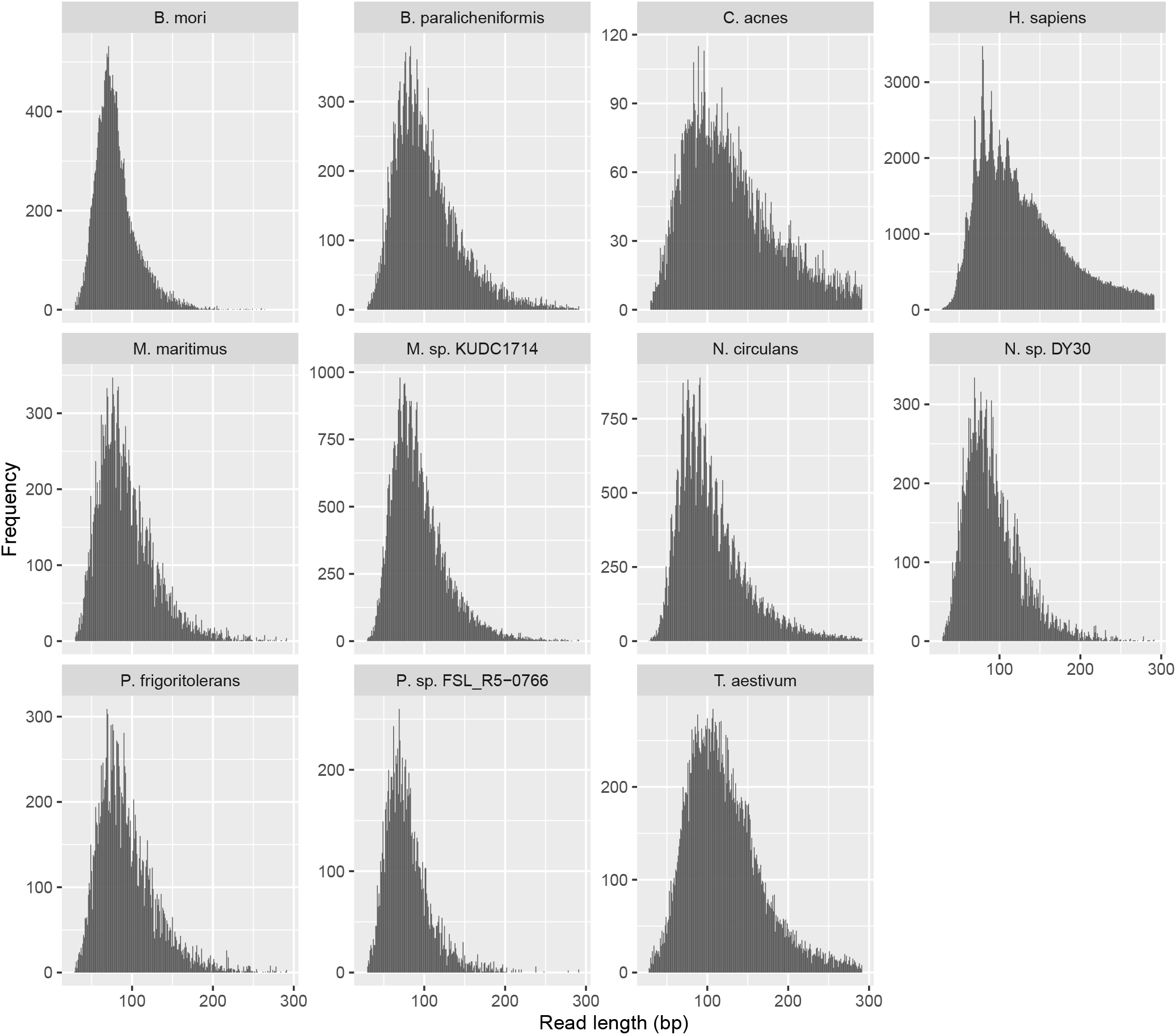
Length distributions of the reads aligned on selected species genomes.

None of the genes used in the literature for the phylogenetic analysis of *Triticum* could be reconstructed from the reads. Therefore, the longest contig (1241 bp, GenBank: PV405787) with high similarity to Poales was used for this purpose. From the BLAST matches of this sequence against the reference genomes of the genera *Hordeum, Triticum* were the following. It showed sequence identity of 71.1-72.6% with chromosome 1H, 2H, 3H, 4H, 5H, 6H, 7H parts of the genome of *Hordeum vulgare* subsp. *vulgare* (CAJHDD000000000.1). While the sequence identity to chromosome fragments 1A, 1B, 2A, 2B, 3A, 3B, 4A, 4B, 5A, 5B, 6A, 6B, 7A, 7B of the *Triticum aestivum* genome (IWGSC CS RefSeq v2.1, JAGHKL00000000000.1) was between 98.2 and 99.1%. The contigs overlapping the PV405787 sequence, generated from reads of the H_vulgare, Ta_PI345242, Tm_TA759, Tt_TU686, Tt_TU692, and Tt_TU1128 samples, were not included in the multiple sequence alignment because they contained more than 5% missing sites. The results of the phylogenetic analysis (with the best substitution model TVM) based on the sequence PV405787 classifiable in the genus *Triticum* are shown in Figure 6.

**Figure 6.**
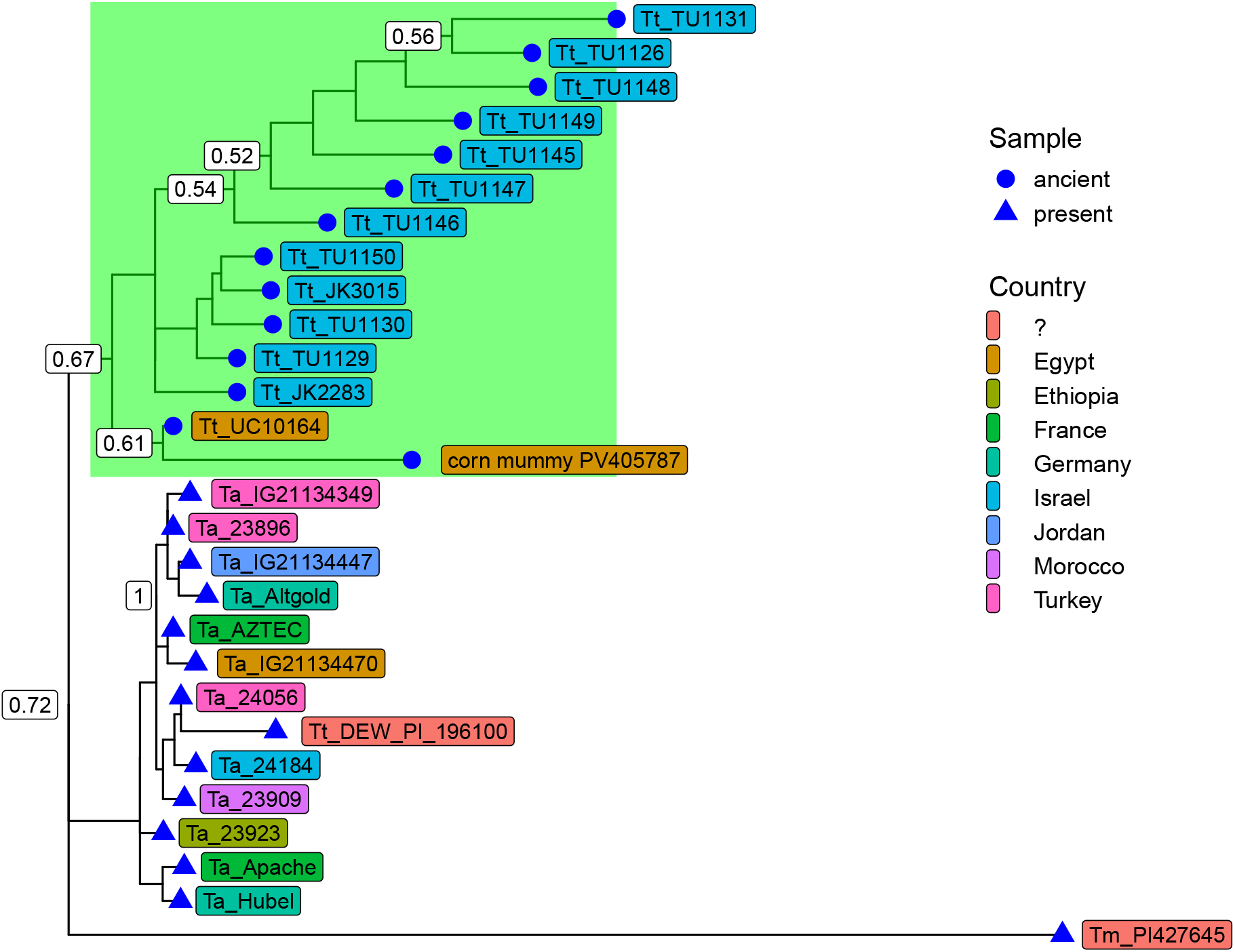
Gene-tree based on the longest wheat origin contig (PV405787) from the corn mummy and the SRA downloaded external samples. Where Ta is *T. aestivum*, Tm is *T. monococcum*, Tt is *T. turgidum* ssp. *dicoccum*. The outer group was the sequence Tm_PI427645. The green band indicates the group containing the corn mummy sequence. Numbers at branches indicate bootstrap support levels (100 replicates). The detailed information of the used samples is summarized in Table 2.

After aligning cleaned reads to the human genome, the average coverage across the 22 chromosomes (excluding Y and MT) was 0.81% (SD: 0.09), with a depth of 0.017 (SD: 0.002). Alignment to the RSRS sequence resulted in 349 reads with 82.02% coverage and 2.56× depth. The coverage distribution of the mitogenome positions was as follows: 53.79% ≥2x, 28.38% ≥3x, 13.59% ≥4x, 5.90% ≥5x, 1.33% ≥6x, 0.40% ≥7x. Based on Haplocheck analysis, mtDNA did not contain any detectable contamination. According to the insufficient depth of mtDNA alignments, schmutzi did not give valuable results for checking contamination. ContamMix estimated mtDNA contamination was 0.79%. The top 5 Haplogrep3 hits of the consensus sequence obtained after deduplication are summarised in Table 3. X-chromosome-based analysis estimated a contamination level of 0.007%. From the ratio of reads matching the X and Y chromosomes, an XY karyotype (*R*_*y*_ = 0.099, 95% CI: 0.093-0.104) was assigned. Aligning the read to the Y chromosome, a coverage of 0.62% and an average depth of 0.01 was obtained, and a haplotype was predicted as J1a2a1a1a~ (path with SNPs: BT: M9087; CT: M5801; J1a2a1a1a : ZS11104A) and E1a2b1a2 ∼ (path with SNPs: BT: M9257, PF1096; CT: M5739, M5801; E: Z15668; E1a2b1a2: M9119, Z14984BT) by pathPhynder and yHaplo, respectively.

**Table 3.** Top 5 haplogroup assignments of RSRS-based mtDNA consensus sequence analysis with Haplogrep3 (metric: Kulczyński, PhyloTree Build: 17). The coverage of the polymorphism positions is 1x: 146, 150, 2885, 4104, 7146, 7256, 7861, 10688, 10915, 13650, 16230, 16278; 2x: 7521, 10810, 10819, 13506, 15301, 16311; 3x: 1018, 2758, 11914, 15944; 4x: 4312, 8655; 5x: 8468, 16189; 6x: 16129, 16187.

| Haplogroup | Quality | Detected polys | Undetected polys |
| --- | --- | --- | --- |
| L3 | 0.863 | 146T 182C! 1018G 2758G 2885T 4104A 4312C 7146A 7256C 7521G 8468C 8655C 10688G 10810T 10915T 11914G 13506C 13650C 15301A 16129G 16187C 16189T 16230A 16278C 16311T |  |
| L3c'd | 0.863 | 146T 152T! 182C! 1018G 2758G 2885T 4104A 4312C 7146A 7256C 7521G 8468C 8655C 10688G 10810T 10915T 11914G 13105G! 13506C 13650C 15301A 16129G 16187C 16189T 16230A 16278C 16311T |  |
| L3h | 0.850 | 146T 182C! 1018G 2758G 2885T 4104A 4312C 7146A 7256C 7521G 8468C 8655C 10688G 10810T 10915T 11914G 13506C 13650C 15301A 16129G 16187C 16189T 16230A 16278C 16311T | 7861C |
| L3b'f | 0.847 | 146T 182C! 1018G 2758G 2885T 4104A 4312C 7146A 7256C 7521G 8468C 8655C 10688G 10810T 10915T 11914G 13506C 13650C 15301A 16129G 16187C 16189T 16230A 16278C 16311T | 15944d |
| L3e'i'k'x | 0.841 | 146T 182C! 1018G 2758G 2885T 4104A 4312C 7146A 7256C 7521G 8468C 8655C 10688G 10810T 10915T 11914G 13506C 13650C 15301A 16129G 16187C 16189T 16230A 16278C 16311T | 150T 10819G |

## Discussion

To the best of our knowledge, no results from the study of the cereal mummy metagenome have been reported to date. Therefore, it is impossible to compare our results with those of others. The key question in interpreting our study’s results is whether our reads, generated in this study, match archaic genomes or are derived from modern contamination.

Due to the unique nature of the sample, DNA extraction has more pitfalls and significantly lower yields than even the usual archaic samples. Prior to our metagenomic analysis, we assumed that a large proportion of the corn mummy sample consisted of plant remains, so we selected an aDNA extraction approach focused on plants. It is possible that higher-quality aDNA could have been obtained by using methods optimized for extracting highly degraded human and environmental DNA. However, we had a limited sample quantity, and our first priority was protecting the corn mummy. Thus, we were unable to perform detailed comparative tests with the aforementioned DNA extraction methods.

The contigs generated from our reads are short, and their number is small compared to the number of reads. The degradation plots based on the reads realigned onto the contigs of Sample 2 show, although not perfectly, patterns that might support a historical origin. An argument in favor of this could be the lack of decay signatures in Sample 1. The trend of the damage plot shows that the G>A transformation increases towards the 3p end of the sequences. At the 5p end, although not as clearly, the C>T transformation also shows an increase. The library-building approach with the Ultra II kit, which uses Phusion-Taq polymerase, is sensitive to uracil and may be responsible for the higher frequency of G>A than C>T. The low number of reads aligned to the contigs (Sample 2: 101,330 reads) may explain the irregular curve pattern. The increase in G>A at the 5p end and C>T at the 3p end is similar to Skoglund et al.^39^, which is not USER-treated. The abrupt decrease in the increasing frequency of C>T at the 5p end and G>A at the 3p end in the last positions resembles the pattern resulting from the internal barcode ligation bias.^40^ The pattern is similar when reads were aligned to selected genomes.

It is reasonable to assume that the corn mummy contains one or more Poales species, and it is very unlikely that the sample is contaminated with fragmented modern genomes from those species. Furthermore, phylogenetic analysis indicates that the sequence from our sample belongs to a group of analyzed external samples containing all archaic *Triticum* sequences. Based on these results, the degradation plot pattern of *T. aestivum* may provide a guide for interpreting the age of the genome fragments of other species. If this is accepted, the archaicity of the *H. sapiens* sequences can be reasonably assumed, as in the case of *B. mori*. The same can be said for the bacterial species, except *C. acnes*, which is more likely to be a modern contaminant.

For the bacteria found in Sample 2, for which DNA damage could be tested and showed a historical pattern of *B. paralicheniformis*,^41^ *Metabacillus* sp. KUDC1714,^42^ *N. circulans*,^43^ their occurrence in the soil is supported by the literature. Although no literature data is available on the occurrence of *Neobacillus* sp. DY30, the origin of the genome we used, is given as rhizosphere soil in the NCBI database. The presence of *P. frigoritolerans* in the rhizosphere has a beneficial effect on plant growth,^44^, which has been explicitly shown for *T. aestivum*.^45^ Although no data are available on *Paenibacillus* sp. FSL_R5-0766 in soil, it is generally acknowledged that numerous *Paenibacillus* species benefit plant growth in soil.^46^ The only bacterium for which no literature data were found on occurrence in soil and rhizosphere was *M. maritimus*. This species is characteristically found in marine sediment,^47^ but other species of the genus occur in sandy soil.^48^

The presence of Poales sequences is not surprising as there is literature data on both *Triticum* and *Hordeum* content in corn mummies.^8,9^ A previous archaeobotanical analysis of the same corn mummy we studied identified barley as a component of the corn.^11^ In our results, more sequences were identified for *Triticum* with minimal presence of *Hordeum*. These two results suggest that it may have been used as a mixed grain for rituals. Although we were unable to use the genes^49^ commonly used for *Triticum* phylogenetic analysis, based on the longest contig (PV405787), our sample, most closely resembles ancient Egyptian and southern Levantine samples.

The interpretation of the *Bombyx* results might be linked to the *Niallia* findings, as *N. circulans* is known to be present in the gut of *B. mori*, among other environments.^50^ The DNA damage of the reads matched to two species suggests that they are likely historical. The larvae of *Bombyx* and its related species feed on the leaves of various plants and pupate either on the plant, on the soil surface, or by digging into the soil. There might be two ways in which a developmental form of the moth could have been introduced into Sample 2. One is that a larva (or pupae) was introduced into the corn mummy with the soil used to create the corn mummy. The other one is that the larva was placed on the open surface of the corn mummy before mummification. There is little probability of the larva entering the aged corn mummy through the fractured surface of the already broken corn mummy and digging into it. This latter case could have happened several decades ago, raising the question of whether the larva would have chosen the old corn mummy for pupation. A further issue with this opportunity is that no related species has been described in museum conditions. If the sequences are indeed from *B. mori*, we need to investigate whether this species may have come into contact with the mummy during its creation. As we may know, silk was not introduced into Egypt until the Ptolemaic period (332-30 BC),^51^ but it was possible at the time of the mummy’s creation. Although the export of *B. mori* moths, larvae or pupae from China was banned until the end of the Middle Ages, if silk had appeared in Egypt, the moth might have had a chance, and in this way *B. mori* might have the opportunity to be incorporated into the mummy. However, it should also be considered that the sample contains not *B. mori* genome fragments but those of a related moth species that occurred in the environment where the mummy was made. Several species of Bombycoidea/Sphingidae are known to be native to Egypt: *Acherontia atropos, Agrius convolvuli, Daphnis nerii, Hippotion celerio, Hyles euphorbiae, Hyles lineata, Hyles livornica, Hyles tithymali deserticola, Macroglossum stellatarum, Theretra alecto, Theretra oldenlandiae*. The NCBI NT database contains complete genomes from these species only for the *H. euphorbiae* and the *H. lineata*. It is also possible that a related moth lived in the area when the mummy was made, but later became extinct, so we don’t even have a trace of its genome. On this basis, we cannot rule out the possibility that the sequences we identified as *B. mori* originated from a related Bombycidae species and lived in Africa.^52^

The haplogrouping of the human sequences from Sample 2 suggests that both mtDNA and the Y chromosome might be of African origin. These haplogroups may have been present in both modern and ancient Egypt. Within the L3 mitochondrial lineage, we found that the L3b, L3d, L3e, and L3f subclades are the most common in Central and West Africa.^53^. L3c is very rare, as reported by Soares et al., who report one sample from East Africa and one from the Middle East.^53^ Rishishwar and King cite rishishwar2017implications, in their global survey of MT haplotype groups, report the distribution by continent based on 2,534 samples. Of these, 272 were L3 haplotypes, of which 74.6% (n=203) were from African populations, 0.37% (n=1) from European populations, 0.74% (n=2) from Indian populations, and 24.3% (n=66) from American populations. However, this latter figure should be supplemented by the fact that 50 of the L3 US samples had known African ancestry (African Caribbean in Barbados, n=26, African ancestry in the South West US, n=24). None of the 504 East Asian samples included in the study belonged to the L3 haplogroup. DNA damage analysis of the reads aligned to the human genome does not rule out their historical origin. In Hallast et al.^54^, the global prevalence of Y-haplotypes was investigated using 1,208 samples. Of these, 150 samples belonged to the E-haplogroup, of which 83.3% (n=125) were from Africa, 8% (n=12) from Central/West Asia, 6% (n=9), 2% (n=3) from South Asia, and 0.7% (n=1) from North Asia. E2 lineage included eight samples of exclusively African origin, including 2 E2a ∼ (M41) samples from Kenya, one E2a∼ from Sudan, one E2b1a (M85) from Gambia, one E2b1a1 (M200) sample each from Congo, Gambia, and Kenya, and one E2b1a1 ∼ (CTS155) sample from Angola.^54^ While the E-haplogroup identified by yHaplo indicates primarily African origin, the type J1a2a1a1a identified by pathPhynder overlaps only partially with this region.^55^ On the other hand, the J1a2a1a haplotype is also found in Egypt, in addition to a higher frequency in the Arabian Peninsula, southern Levant, and southern Iraq.^55^ Keita’s review of the diversity of Y chromosome variants supports our findings.^56^

Consistent with our findings, others have identified human remains in European collections of Egyptian archaeological materials, including canopic jars.^57,58^ The authors of these studies emphasize that non-historical contamination cannot be ruled out in these cases, a point we agree with based on our own results. Similar to our results, Rayo et al.^58^ identified human mtDNA haplogroups from canopic jar samples, the level of degradation of which casts doubt on their ancient origin. The haplogroups they identified, similar to ours, occur in the present-day African population, supporting the possibility of a modern origin. As in the studies mentioned above, we also have some uncertainty about the age of the DNA examined, but there are additional considerations.

If Sample 2 had been taken from the mummy’s surface, one might think that DNA sequences belonging to the mentioned haplogroups could have remained there from handling in Egypt during the modern period. Nevertheless, as this sample was taken from a depth of a few millimeters, it is difficult to imagine that human DNA would be present due to human touch. But, it is possible that some human DNA present in any human DNA-containing fluid may have infiltrated the mummy and been found. For instance, this could have happened if the crypt in which the mummy was kept had been contaminated with human sewage, which could have soaked the mummy. As this case could have occurred before the mummy was placed in the museum, the DNA damage that has occurred in the decades since could have produced the smiley plot patterns we have seen. In addition to these possibilities, the possibility that human DNA was introduced during the mummification process cannot be completely ruled out. Since the mummy is described as having been molded, it was obviously made by human hands, so the genetic material from shed epithelial cells may have been preserved in a lucky case. At the same time, it cannot be ruled out that human DNA may have come from human feces (e.g., night soil), which may have been mixed with the seed/soil/water mixture to promote grain growth. It is worth noting that the mummy was dried in the sun during its preparation, creating a dry environment that helped preserve the DNA. Although others have reported results based on similarly fragmented data (e.g., Ziesemer et al.^59^ used 3–97× depth samples, with coverage between 62–97%), the reliability of our haplotyping results would be improved by more complete sequences.

## Conclusion

A metagenomic analysis of the corn mummy offers an opportunity to better understand its components. However, even with the latest genetic dating methods, interpreting the results requires caution. Even though the samples taken from the corn mummy are likely to contain both modern and ancient genetic material, sequences that are more likely to be from Egypt in the 3rd century BC were identified. While certain bacterial sequences appear to be of modern origin, sequences from an ancient wheat cultivar, a silkmoth-related insect with its gut bacteria, and an African human male.

## Declarations

## Acknowledgements

In memory of Vendel Dolmány O.Cist. We want to thank Lilla Füleki, a colleague of the National Museum Conservation and Storage Centre (OMRRK) – Museum of Fine Arts, Budapest, Hungary, for her enthusiastic suggestions and support. We are also grateful to Shomarka Omar Yahya Keita for his help clarifying our wording and putting the results into context.

## Author contribution statement

NS takes responsibility for the data’s integrity and the data analysis’s accuracy. DM, IC, and NS conceived the concept of the study. AGT, FJK, KK, and SÁN got the samples from the corn mummy. BP and GM did the DNA extraction and NGS sequencing. NS performed the bioinformatic analysis and visualization. AGT, DM, NS, and SÁN participated in the drafting of the manuscript. AGT, BP, DM, FJK, GM, IC, KK, NS, and SÁN critically revised the manuscript for important intellectual content. All authors read and approved the final manuscript.

## Data Availability

The raw short read data are publicly available and accessible through the PRJNA1128416 (Sample 1: SAMN42050348; Sample 2: SAMN42050349) from the NCBI Sequence Read Archive (SRA).

Reviewer link: https://dataview.ncbi.nlm.nih.gov/object/PRJNA1128416?reviewer=ve2k8luao0da4qtkasur1a4icv Upon reasonable request, the corresponding authors of this article will provide unrestricted access to the original data and code.

## Ethics approval and consent to participate

Not applicable.

